# A transcriptomics-native foundation model for universal cell representation and virtual cell synthesis

**DOI:** 10.64898/2026.04.12.718016

**Authors:** Xiaohui Jiang, Jichun Xie

**Affiliations:** Department of Biostatistics and Bioinformatics, Duke University, Durham, NC, USA; Department of Mathematics, Duke University, Durham, NC, USA; Department of Computer Science, Duke University, Durham, NC, USA

**Keywords:** single-cell omics, spatial omics, generative pre-training, transcriptomics-native foundation model, virtual cell synthesis

## Abstract

Current single-cell foundation models rely on language-model architectures that ignore transcriptomic data distributions, often underperforming specialized methods. We introduce xVERSE, a transcriptomics-native foundation model coupling batch-invariant representation learning with the probabilistic generation of expression profiles. xVERSE outperforms the leading foundation and batch-effect correction methods in representation learning by **17.9%** and **11.4%**, respectively, successfully preserving biological heterogeneity while diminishing batch effects. Furthermore, xVERSE surpasses the second-best spatial imputation method by **34.3%** and uniquely synthesizes virtual cells indistinguishable from biological data (AUROC **≈ 0.5**). As a powerful data-augmentation engine, xVERSE utilizes these high-fidelity virtual cells to enable accurate clustering and marker detection in tiny datasets—resolving rare cell types with as few as four cells—while improving the generalizability of cross-modality predictions across diverse pathological states. These results establish xVERSE as a transformative framework unlocking analytical capabilities beyond conventional models.

## 1 Introduction

The rapid expansion of single-cell and spatial sequencing has generated massive population-level atlases charting cellular profiles across tissues, developmental stages, and diseases [1–12]. These resources present a unique opportunity to learn universal cell and gene representations and generative transcriptomic distributions. Universal cell and gene representations are essential for integrating heterogeneous datasets to uncover biological consensus across studies, gene panels, and technologies [13–23]. Concurrently, cell transcriptomic generative modeling enables the accurate imputation of unmeasured genes and the synthesis of high-fidelity virtual cell, augmenting downstream analysis on small datasets and other machine-learning model training [24–30].

Several transcriptomics foundation models have recently emerged, adapting large language model architectures—such as BERT [31] and GPT [32]—to single-cell data (e.g., Nicheformer, scGPT, Geneformer) [33–39]. While some of these models, such as scGPT, employ generative pre-training objectives, their primary goal is to predict masked tokens to learn robust cell and gene representations, rather than to explicitly model the full probability distributions of the transcriptome. Consequently, they function primarily as representation learners rather than generative engines capable of synthesizing high-fidelity virtual cells. Furthermore, despite their significant computational demands, these models often do not outperform some traditional mono-task methods in batch-effect correction, unmeasured gene imputation, or inference from small datasets[40–46]. This limitation likely stems from a fundamental architectural mismatch: the sequential priors inherent to language models are ill-suited for the unordered, high-dimensional, and sparse transcriptomic data [47–50], restricting their ability to capture the transcriptomic distributional characteristics required for high-fidelity generation and robust downstream inference.

To address these limitations, we introduce xVERSE, a deep generative foundation model that unifies universal representation learning with probabilistic transcriptomic reconstruction. By training on large-scale single-cell and spatial transcriptomic data, xVERSE learns to disentangle intrinsic biological signals from technical confounders, such as batch and platform effects, yielding robust and panel-invariant representations [51–53]. Unlike existing foundation models, xVERSE directly models cellular genes’ joint probability distributions. This allows it to output a generative transcriptomic distribution for any given cell embedding—derived from either full or partial gene panels—enabling the imputation of unmeasured genes, the synthesis of virtual cells, and the augmentation of small datasets to uncover weak biological signals.

xVERSE incorporates four key architectural innovations tailored to transcriptomic data. First, instead of imposing an artificial sequential structure as done in previous Transformer-based models [33–39], xVERSE directly models the raw gene counts, respecting the native unordered and high-dimensional nature of transcriptomic profiles [47–50]. Second, it employs a panel-aware stochastic gene-masking strategy for robust self-supervised learning across varying gene panels [54–59]. Third, it utilizes a Gradient Reversal Layer (GRL) to explicitly separate biological variation from technical confounders, ensuring the learned representations are invariant to batch and technology differences[60]. Finally, it models the underlying data-generating process using cell- and gene-specific Poisson distributions, guided by a distribution reconstruction loss to ensure the generation of biologically realistic cell profiles.

In this paper, we demonstrate the power of xVERSE across three key domains (Fig. 1a).

**Fig. 1.**
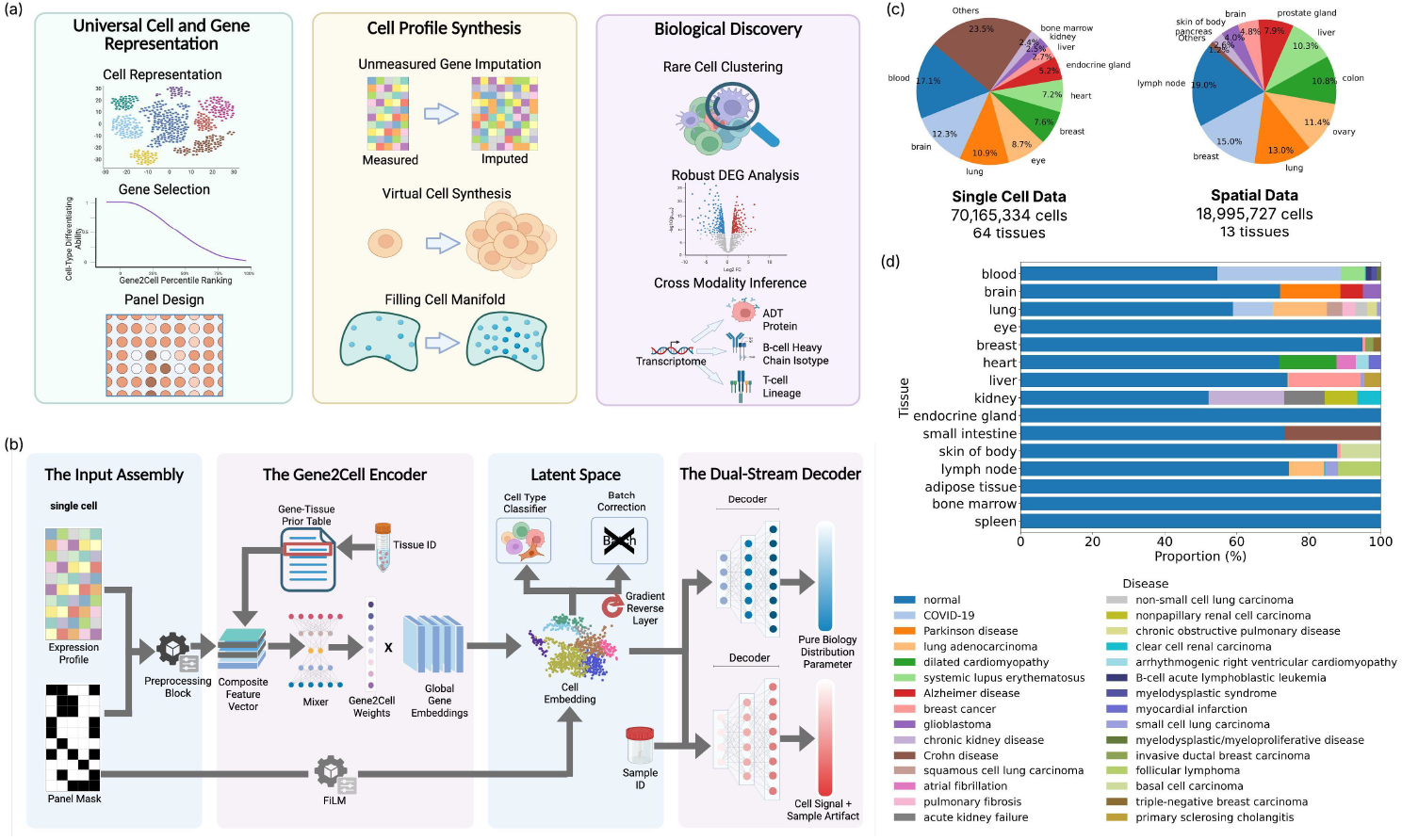
Overview of xVERSE: A transcriptomics-native foundation model. **a**, xVERSE’s power in three key domains: (1) Universal cell and gene representation; (2) Cell profile synthesis; and (3) Biological discovery. **b**, The xVERSE architecture unifies representation learning and generative modeling. It features a Gene2Cell Encoder that projects expression profiles into a batch-invariant latent space, and a Dual-Stream Decoder that reconstructs cell-specific signals and technical artifacts separately. **c**, Tissue composition of all cells in the training dataset, grouped by single-cell (64 tissues, 70M cells) and spatial transcriptomics (13 tissues, 19M cells) platforms. Complete statistics are provided in Suppl. Table 1. **d**, Disease composition across the top 15 tissues by cell count, showing the diversity of disease states in the training data. Complete statistics are provided in Suppl. Table 2.

- **Universal Cell and Gene Representation**. xVERSE learns robust, cell-type-aware representations that are invariant to batch effects, gene panel variations, and technological differences. Additionally, it derives informative gene representations and gene-specific Gene2Cell scores: Top-ranked genes that enhance the signal-to-noise ratio for resolving cellular heterogeneity, and thus can be prioritized for panel design.
- **Cell Profile Synthesis**. xVERSE is the first foundation model capable of synthesizing “virtual cells” that are transcriptomically indistinguishable from biological cells (AUROC ≈ 0.5 in classification). Synthesizing high-fidelity virtual cells can fill in the gap of rare cell types in the observed cell manifold and improve downstream analysis performance. This generative capability also enables the precise imputation of unmeasured genes in spatial transcriptomics without relying on external references. Head-to-head comparisons show xVERSE significantly outperforms traditional task-specific imputation methods, even in zero-shot settings.
- **Biological Discovery**. xVERSE leverages its generative capabilities to accelerate discovery. In the context of small datasets, it improves rare cell type clustering and enables robust differential expression gene (DEG) analysis. Furthermore, in a challenging cross-modality prediction benchmark, xVERSE augmentation significantly improved the training of downstream machine learning models. Notably, models augmented with xVERSE-synthesized virtual cells demonstrated superior generalization to unseen pathological states, transferring knowledge from normal recovery states to patients with cardiac allograft vasculopathy and non-specific graft dysfunction.
- These applications exemplify xVERSE’s role as an integrative tool for accelerating biological discovery, offering potential that extends beyond these examples to deepen our understanding of complex biological mechanisms.

## 2 Results

### 2.1 xVERSE Model Overview

xVERSE is a generative foundation model that unifies representation learning and whole-transcriptome expression distribution learning across diverse single-cell and spatial platforms (Fig. 1b). Uniquely, xVERSE emphasizes the probabilistic modeling of the entire transcriptome. By decoding latent representations, the model generates cell- and gene-specific Poisson distributions that characterize the underlying expression probability for every gene. This capability allows xVERSE to synthesize virtual cells—generating high-fidelity *in silico* transcriptomes based on biological cell templates—thereby capturing the full complexity of gene expression distributions.

Complementing this generative capacity, xVERSE also projects inputs into a batch-invariant latent biological embedding space, enabling robust cell-type inference and data integration.

To train xVERSE, we curated a large-scale pan-tissue pan-disease cross-technology single-cell and spatial transcriptomics database, comprising over 89 million cellular profiles (Fig. 1c). Among them, over 70 million cells were profiled using high-throughput droplet-based single-cell or single-nucleus RNA-sequencing technologies [1], spanning 64 distinct tissues and 138 disease states (including normal) (Fig. 1c,d; Suppl. Tables 1 and 2); about 19 million cells were profiled using *in situ* imaging-based spatial transcriptomics technologies [61] including Xenium v1, Xenium Prime [62], MERFISH [63], and CosMx [64], spanning 13 tissues (Fig. 1c; Suppl. Table 1). This comprehensive dataset enables xVERSE to learn robust universal cell representations and cell-gene-specific expression distributions.

### 2.2 xVERSE Learns Robust Universal Cell Representations Across Studies and Panels

Robust universal cell representations are critical for integrating fragmented datasets into a cohesive biological atlas. We benchmarked the zero-shot performance of xVERSE against three leading transcriptomics foundation models (scGPT, Nicheformer, and Geneformer) using two independent datasets: a human healthy liver scRNA-seq atlas [65] and a human healthy and amyotrophic lateral sclerosis (ALS) motor cortex scRNA-seq atlas [66] (Fig. 2a, b). To ensure no data leakage, we selected two datasets published after our pre-training data collection. For rigorous and fair comparison, all models were applied directly without fine-tuning. xVERSE consistently achieved the highest cell-type Average Silhouette Width (ASW) scores [67] across both datasets, outperforming the second-best foundation model, scGPT [38], by 17.9% on average, demonstrating superiority in capturing cell heterogeneity. Notably, xVERSE maintained its superiority even when inputs were restricted to limited gene subsets matching spatial technologies (e.g., the 5,000-gene Xenium Prime and 377 or 266-gene tissue-specific panels [68]), where it surpassed scGPT by 19.4% and 15.8% respectively, demonstrating its consistent superiority across varying data modalities and gene panels. Furthermore, xVERSE has a high computation efficiency, with an average inference speed 59.7% faster than the second-best model, scGPT (Fig. 2c). Also, we evaluated the computational efficiency of all foundation models relative to its inference time (Suppl. Fig. 1). Across different tissues and gene panel sizes, xVERSE consistently achieves the highest ASW scores while maintaining the lowest inference time.

**Fig. 2.**
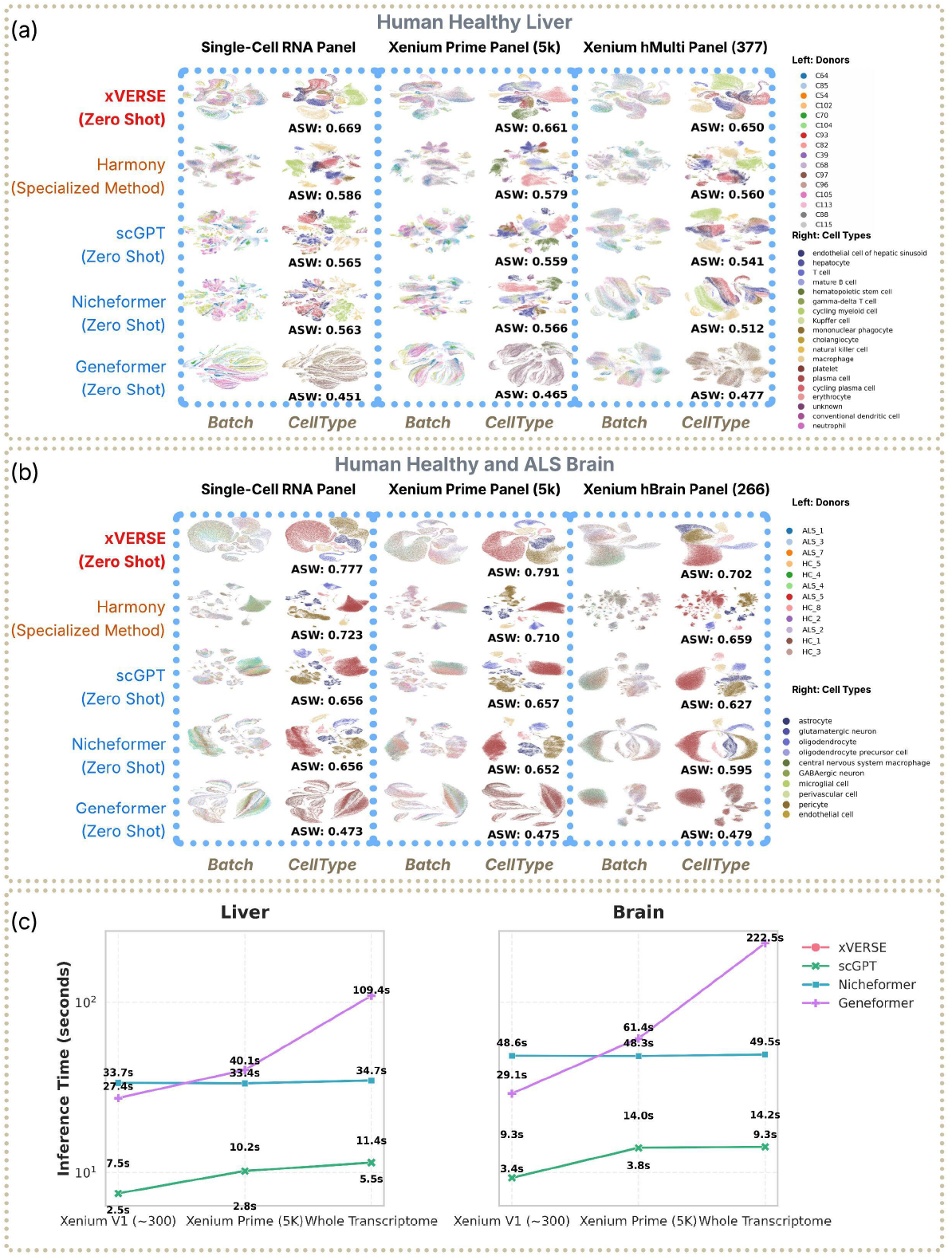
**a)-b) Cell projections to the representation spaces constructed by xVERSE, other leading foundation model (scGPT, Nicheformer, and Geneformer) without fine-tuning, and dataset-specific method Harmony.** a) Human healthy liver scRNA-seq atlas; b) Human healthy and amyotrophic lateral sclerosis (ALS) motor cortex scRNA-seq atlas. Full transcriptomic and restricted gene sets matching Xenium Prime (5K genes) and Xenium tissue-specific panels (liver: 377 genes; brain: 266 genes) were used to profile cells, respectively. Cells are colored by batch id (left panel) and cell-type annotations (right panel) provide by the datasets. **c) Zero-shot inference time comparison between xVERSE and other leading foundation models (scGPT, Nicheformer, and Geneformer)**. Zero-shot inference time (avaraged over all samples) is shown in seconds (log scale).

Next, we benchmarked xVERSE against Harmony [19], a widely used task-specific batch-correction method for single-cell data. Remarkably, in zero-shot inference, xVERSE outperformed Harmony by achieving a consistent increase in cell-type ASW (+11.4% on average), while other leading foundation models underperformed Harmony. This result challenges the prevailing view that general-purpose foundation models often underperform task-specific tools in zero-shot mode [40–46]. By explicitly disentangling biological variation from technical confounders via its architectural design, xVERSE delivers a “best-of-both-worlds” solution: the scalability and universality of a foundation model with the precision and robustness of a specialized integration tool.

### 2.3 Gene2Cell scores enable biological interpretability and guide panel design for spatial transcriptomics

Beyond representation learning and generation, xVERSE provides intrinsic biological interpretability through a “Gene2Cell Score.” This score quantifies the contribution of each gene to the learned cellular embedding for every individual cell, effectively identifying marker genes that drive each cell’s biological identity.

To validate the biological significance of these scores, we examined their capability to capture cell heterogeneity. For each of five different cell cohorts, we stratified genes into quartiles based on their average Gene2Cell scores, and then project cells with only genes in each quartile to the embedding space, and evaluated whether each quartile gene set can separate cell types quantified by cell-type ASW score (Fig. 3a). Strikingly, embeddings derived solely from the top 25% high-scoring genes have higher ASW scores than those using the full gene panel, consistently across all five cell cohorts. Other quantile-level stratification show similar results (Suppl. Fig. 2). This indicates that the Gene2Cell score successfully filters out noise and prioritizes the most biologically relevant genes for defining cell types.

**Fig. 3.**
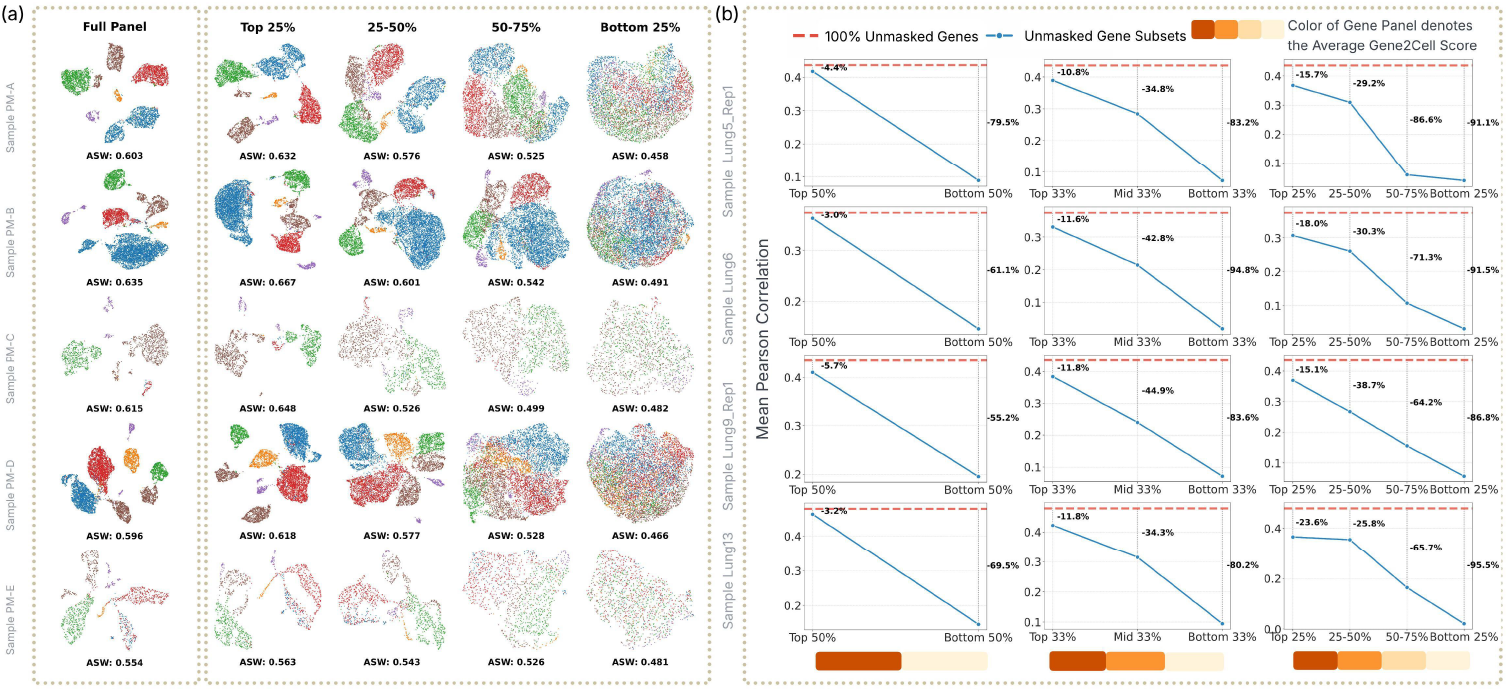
Gene2Cell scores enable biological interpretability and guide panel design. **a**, Impact of Gene2Cell score-based gene selection on single-cell representation quality. Genes were stratified into quartiles based on their Gene2Cell scores, and cell embeddings were evaluated using ASW. Embeddings derived from the top 25% of high-scoring genes consistently yielded higher ASW scores than those using the full gene panel, demonstrating that Gene2Cell scores effectively filter noise and prioritize biologically relevant features. **b**, Evaluation of imputation performance using gene subsets selected by Gene2Cell scores in spatial transcriptomics. Using only the top 50% of high-scoring genes resulted in negligible accuracy loss compared to the full panel, validating the utility of Gene2Cell scores for guiding probe selection in targeted spatial technologies.

We further demonstrated the practical utility of this scoring in spatial transcriptomics. We evaluated imputation performance using different subsets of genes stratified by their average Gene2Cell scores (Fig. 3b). Remarkably, using high-scoring genes, such as the top 50% high-ranking genes, resulted in only a minor drop in imputation accuracy compared to using the full panel excluding the need-to-be-imputed genes. In contrast, with lowly ranked genes, such as bottom 50%, imputation accuracy could drop by more than 79%. The results suggest that xVERSE’s Gene2Cell scores can effectively guide gene panel optimization—an essential step for many spatial transcriptomics technologies—to recover maximal biological information with a minimized gene panel [54–59].

### 2.4 xVERSE synthesizes high-fidelity virtual cell transcriptomic profiles

xVERSE represents an innovative paradigm shift as the first foundation model capable of synthesizing virtual cells from a seed template. To assess its synthetic fidelity, we fine-tuned xVERSE on five distinct single-cell datasets (PM-A to PM-E) [69] and generated three corresponding virtual cohorts for each dataset. We first examined the distribution of gene expression counts, focusing on the top 20 highly variable genes (HVGs). For representative genes with distinct expression profiles—such as *RPLP1, RPS8, GSN, MGP*, and *EEF1A1*—xVERSE accurately reproduced the UMI count distributions of biological cells, faithfully capturing both the characteristic sparsity and dynamic range of real biological data (Fig. 4a). Other top HVGs also exhibited similar patterns, further validating xVERSE’s ability to synthesize high-fidelity virtual cells (Suppl. Figs. 3–7).

**Fig. 4.**
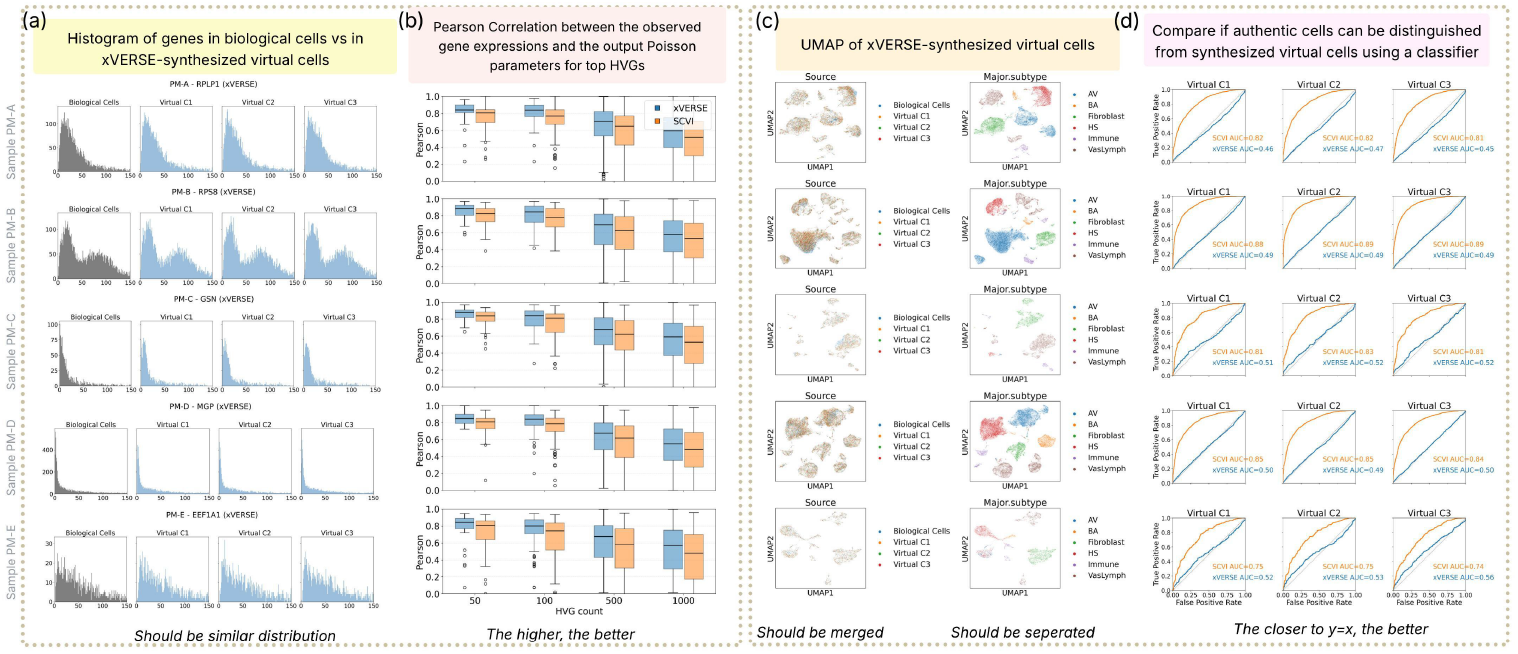
xVERSE synthesizes high-fidelity virtual cells that are distributionally accurate and transcriptomically indistinguishable from biological cells. **a**, Distributional fidelity of gene expression counts. Histograms display UMI count distributions for representative HVGs— including *RPLP1, RPS8, GSN, MGP*, and *EEF1A1*—across five distinct single-cell datasets (PM-A to PM-E). xVERSE-generated cohorts (Virtual Datasets 1–3) accurately reproduce the sparsity and dynamic range of the authentic biological data without mode collapse. **b**, Preservation of gene expression rankings. Boxplots compare the Pearson correlation of HVG rankings between authentic and synthetic profiles generated by xVERSE (blue) versus scVI (orange). Comparisons across varying feature set sizes (50 to 1000 HVGs) demonstrate that xVERSE consistently yields higher correlations, indicating superior retention of relative expression structures. **c**, Conservation of manifold structure and cellular identity. UMAP projections of combined authentic and virtual datasets. **Left:** Cells colored by source (Biological vs. Virtual Datasets 1–3) exhibit seamless integration, confirming the absence of generation-specific artifacts. **Right:** The same embeddings colored by cell type reveal that virtual cells maintain distinct, biologically coherent clusters (e.g., immune cells, fibroblasts) matching the original annotations. **d**, Quantitative assessment of indistinguishability via adversarial classification. ROC curves show the performance of binary classifiers trained to discriminate between authentic and generated cells. Classifiers fail to distinguish xVERSE-synthesized cells (blue curves), yielding AUROC scores near 0.5 (random guessing). In contrast, scVI-synthesized cells (orange curves) are readily detectable (AUROC ≈ 0.75-0.9), highlighting xVERSE’s superior capacity to capture high-dimensional biological realism.

We then benchmarked xVERSE against scVI [25], a leading non-foundation-model-based generative model, by evaluating the preservation of gene expression rankings. We calculated the Pearson correlation of HVG rankings between biological and virtual datasets across varying gene set sizes (50 to 1000 HVGs). xVERSE consistently yielded higher correlation scores than scVI (+9.2% on average), indicating a superior ability to preserve the finer relative expression structures that define cellular identity (Fig. 4b).

Finally, we evaluated whether xVERSE-synthesized virtual cells retained the complex biological heterogeneity of the originals while being indistinguishable. In UMAP [70] projections, xVERSE-synthesized virtual cells integrated seamlessly with biological cells while maintaining distinct cell-type clusters (e.g., immune cells, fibroblasts) with high fidelity (Fig. 4c). To rigorously quantify the virtual cells’ indistinguishability, we trained binary classifiers to discriminate between biological and virtual cells synthesized using three independent runs. Remarkably, the classifiers failed to distinguish xVERSE-synthesized virtual cells from biological cells, yielding Area Under the Receiver Operating Characteristic Curve (AUROC [71]) scores near 0.5 (random guessing) (Fig. 4d). In contrast, cells generated by scVI were easily detected, with AUROC scores consistently exceeding 0.7. These results demonstrate that xVERSE surpasses existing generative models, synthesizing high-fidelity virtual transcriptomic profiles that capture the subtle, complex, and high-dimensional dependencies of real biological systems.

### 2.5 xVERSE imputed unmeasured genes more accurately than specialized methods

Imputing unmeasured genes is a critical task in spatial transcriptomics, as many targeted gene panels cannot include all genes of interest. For instance, widely used Xenium tissue-specific panels typically probe fewer than 500 genes. Although larger panels have been developed, a fundamental trade-off exists in probe-based technologies: increasing the panel size often paradoxically reduces mRNA capture efficiency due to probe competition, saturation, and optical crowding [72–75]. Consequently, researchers often choose smaller, high-sensitivity panels, leaving the vast majority of the transcriptome unmeasured. Accurate imputation of these missing genes is therefore essential to broaden the analytical scope and uncover hidden biological signals within restricted spatial datasets.

To evaluate xVERSE’s imputation performance, we conducted a rigorous “masking” experiment on 4 samples in the CosMx lung NSCLC dataset. Specifically, we uniformly masked the top 50 HVGs and tasked the models with reconstructing their expression profiles. xVERSE successfully recovered the spatial distributions of these masked genes, with imputed expression patterns closely matching the ground truth. This is visually evident in marker genes such as *KRT19, CD74, COL1A1*, and *COL3A1*, where xVERSE faithfully reproduced the complex spatial architectures (Fig. 5a).

**Fig. 5.**
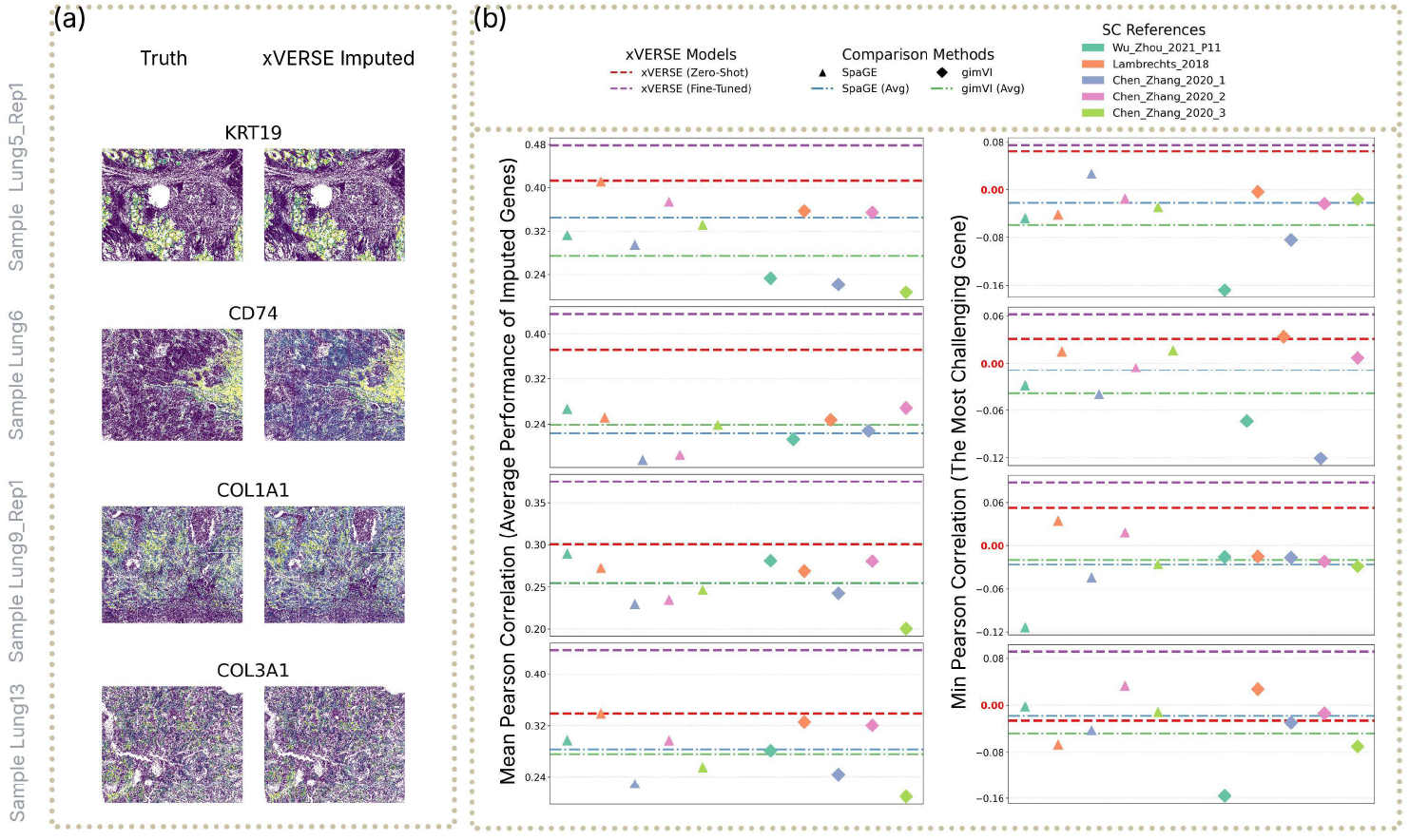
xVERSE accurately imputes unmeasured genes in spatial transcriptomics, out-performing specialized reference-dependent methods. **a**, Spatial reconstruction of masked genes. Comparison of ground truth (left columns) and xVERSE-imputed (right columns) expression maps for representative marker genes—*KRT19, CD74, COL1A1*, and *COL3A1* —across four distinct samples. In this “masking” experiment simulating a targeted panel, xVERSE faithfully recovers complex spatial architectures and expression gradients solely from a restricted subset of input genes. **b**, Quantitative benchmarking against leading specialized methods (SpaGE and gimVI). **Left:** Mean Pearson correlation of the top 50 HVGs between imputed and true profiles. Comparison methods (triangles and diamonds) exhibit high variance depending on the external single-cell reference used (color-coded legend). In contrast, xVERSE (dashed lines; red: zero-shot, purple: fine-tuned) consistently outperforms these methods without requiring any external reference data (zero-shot) or with the same reference data (fine-tuned). **Right:** Robustness stress-test evaluating the minimum Pearson correlation (performance on the single most difficult-to-predict gene among the top 50 HVGs). While specialized methods frequently suffer catastrophic failure (yielding invalid negative correlations), xVERSE demonstrates superior resilience, maintaining positive correlations even in the most challenging prediction scenarios.

We benchmarked xVERSE against two leading spatial imputation methods, SpaGE [29] and gimVI [27], by comparing the Pearson correlations of the top 50 HVGs (masked genes) between imputed and true expression profiles. A critical limitation of these specialized methods is their dependence on external single-cell reference datasets. We found that their performance is highly sensitive to the choice of reference, with imputation accuracy fluctuating wildly. For example, in the Lung5 analysis, gimVI’s accuracy varied from a mean Pearson correlation of 0.2074 to 0.3572, and SpaGE’s performance fluctuated between 0.2948 and 0.4115, merely by changing the reference dataset. In contrast, xVERSE eliminates this reference dependency: on the same sample, the zero-shot xVERSE model achieved a Pearson correlation of 0.4130 without any external reference, matching the best-case performance of SpaGE and outperforming all gimVI runs. Fine-tuning xVERSE on the same references yielded an even higher correlation of 0.4785. A comprehensive evaluation across all samples confirms this general trend: xVERSE consistently delivers superior and stable imputation accuracy, outperforming the second-best specialized method by 34.3% on average, regardless of the reference single-cell data used (Fig. 5b, left panel and Suppl. Fig. 8).

Finally, we tested model robustness in a “stress test” regime, focusing on the gene that was the most difficult to predict among the top 50 HVGs. In this challenging scenario, specialized methods failed catastrophically, yielding invalid negative correlations (Fig. 5b, right panels). Conversely, xVERSE demonstrated remarkable resilience, maintaining positive correlations *>* 0.06 after fine-tuning. While absolute scores in this regime are naturally lower, xVERSE’s ability to avoid catastrophic failure highlights its reliability and stability as a robust solution for whole-transcriptome imputation.

### 2.6 xVERSE amplifies latent biological signals to resolve rare cell populations in low-cell-count datasets

Low-cell-count datasets—such as pilot studies or rare clinical samples—often lack sufficient statistical power to resolve subtle cellular heterogeneity, particularly for low-abundance cell types[76–80]. xVERSE can generate high-fidelity virtual cells that amplify latent biological signals in rare populations, thereby increasing effective sample size and improving the performance of downstream analyses.

First, we assessed the capacity of xVERSE to distinguish between biologically closely related cell types within a low-cell-count imbalanced cell population–the minor population comprise as few as 4–10 cells out of a total of 60. Under this scenario, standard analysis pipelines (e.g., Leiden [81] clustering) failed to differentiate the minor and major cell-type populations, a known challenge in rare cell-type detection. Across varying resolutions (0.5, 1, or 2), Leiden either merged the minor cell type into the major population or fragmented the cells into spurious clusters, consistently yielding Adjusted Rand Index (ARI) scores [82] near zero. However, by applying xVERSE to synthesize 5 virtual cells per biological seed cell and re-evaluating the pipeline, we successfully discovered the rare cell-type population. Applying the same clustering method (Leiden) but following xVERSE’s augmentation significantly improved cell-type separation, in some instances achieving perfect clustering (Fig. 6a).

**Fig. 6.**
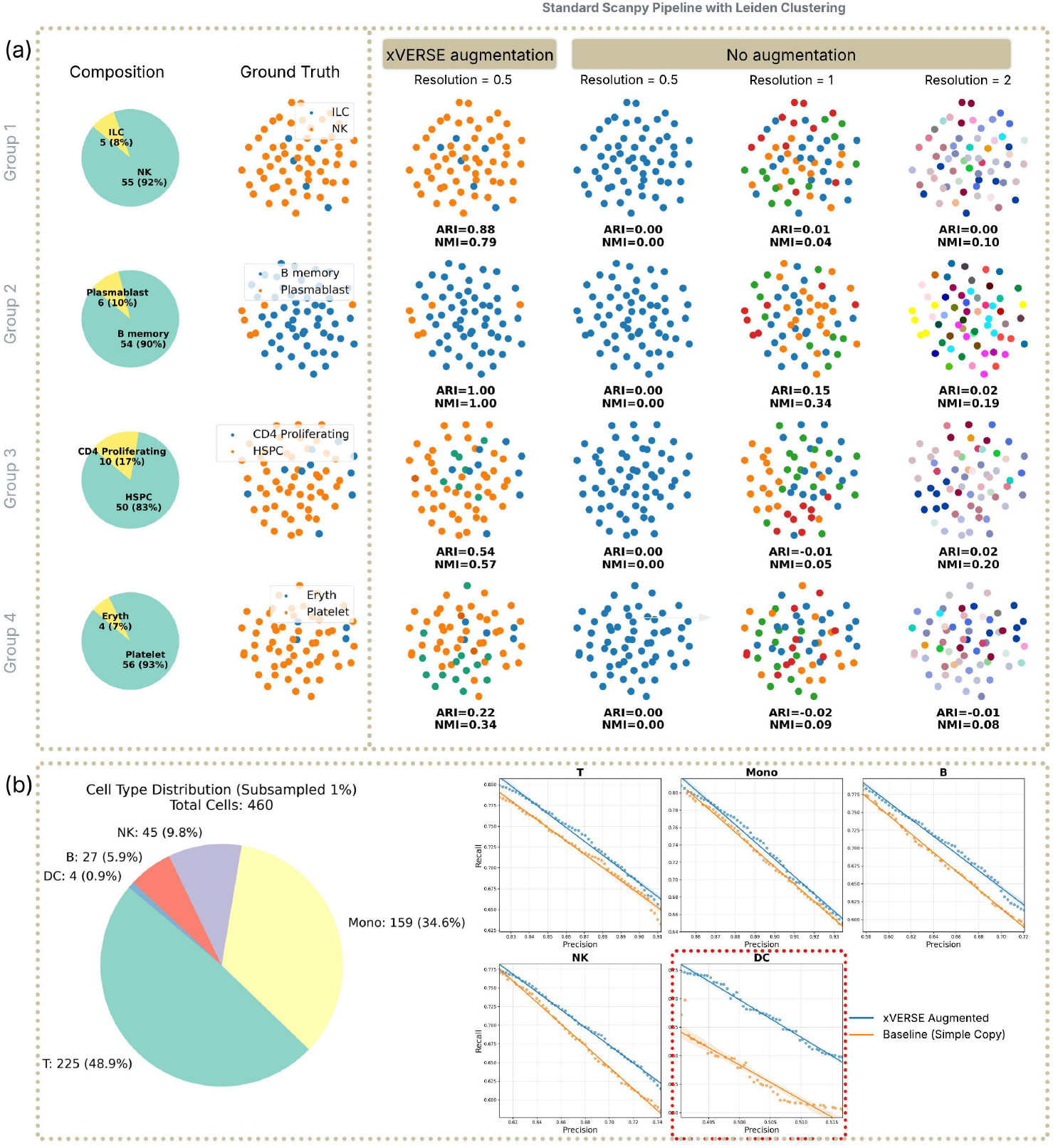
xVERSE amplifies latent biological signals to resolve rare cell populations and rescue statistical power in small-scale datasets. **a**, Evaluation of xVERSE in distinguishing biologically closely related cell types within small-scale, highly imbalanced cell populations (total cells ∼60; minor population comprised of 4–10 cells). The leftmost column (“Composition”) displays the proportion of dominant versus minor cell types for four distinct biological groups. Scatter plots compare the clustering performance of “Ground Truth” labels against its 1% subsampled dataset and 1% subsampled dataset augmented with xVERSE (synthesizing 5 virtual cells per biological seed). Performance is quantified using Adjusted Rand Index (ARI) and Normalized Mutual Information (NMI) [83]. **b**, Assessment of xVERSE’s ability to rescue DEG detection in a subsampling experiment. **Left:** Pie chart showing the cell type distribution of a dataset subsampled to 1% of its original size (*N* = 460), creating an ultra-rare Dendritic Cell (DC) population (*n* = 4; 0.9%). **Right:** Precision-Recall curves benchmarking DEG identification against ground truth derived from the full dataset (*p*-value < 0.05). The performance of xVERSE augmentation (100 virtual cells per seed; blue line) is compared to baseline oversampling (simple replication; orange line) across significance thresholds ranging from *p* = 10^*−*8^ to 0.05.

Next, we evaluated whether xVERSE could improve DEG identification in low-cell-count datasets. We subsampled 1% of an immune cell dataset to retain original cell type proportions, resulting in a dataset containing only 4 (0.9% of the total) dendritic cells (DC). Defining “ground truth” DEGs using the full, original dataset (*p*-value < 0.05), we compared two augmentation strategies on the small dataset: (1) xVERSE augmentation (100 virtual cells synthesized per biological cell) and (2) baseline oversampling (naively copying each cell 100 times). We benchmarked the precision and recall of identified DEGs against the ground truth across a comprehensive range of significance thresholds (*p*-value 10^*−*8^ to 0.05). Consistently, xVERSE augmentation yielded superior performance than the baseline oversampling. Notably, for the ultra-rare DC population, xVERSE augmentation significantly outperformed the baseline oversampling, improving recall by approximately 0.1 at any fixed precision level (Fig. 6b). Collectively, these results confirmed that xVERSE-synthesized virtual cells effectively rescue statistical power and enable deeper biological insights from low-cell-count datasets.

### 2.7 xVERSE’s high-fidelity virtual cell augmentation enhances generalizable machine learning model training

Beyond augmenting data to enhance standard single-cell analysis, xVERSE serves as a potent data augmentation engine capable of enhancing the robustness and accuracy of other machine learning model training [84]. Some context-specific datasets contain cells collected from subjects with specific disease status, age, sex, or other clinical features; models trained on such datasets may fail to generalize to new contexts [49, 85], leading to suboptimal performance [26]. This challenge often arises from biological state shifts of cells between contexts: cells that are under-represented and poorly learned in the training data may become dominant in the test data, where the model fails to fit. To address this, xVERSE synthesizes high-fidelity virtual cells that populate the latent cell-state manifold, thereby boosting signal detection in under-represented populations and improving model generalization.

As an example, we trained and tested cross-modality prediction models on different cohorts: models were trained exclusively on PBMC samples from post-transplant clinically stable (“normal”) individuals but tested on two different pathological patient cohorts—Non-specific Graft Dysfunction (NGD) and Cardiac Allograft Vasculopathy (CAV)—without further fine-tuning. The cross-modality prediction task is using cell transcriptomic profile to predict antibody-derived tag (ADT) protein levels (measured by CITE-seq), B-cell heavy-chain isotype (measured by VDJ-seq), and T-cell lineage (measured by VDJ-seq). As shown in Fig. 7b, the pathological cohorts display pronounced shifts in B-cell heavy-chain isotype usage and highly imbalanced T-cell lineage distributions relative to the training cohort, whereas the z-scored ADT profiles are broadly similar with only modest cohort-specific deviations. Together, these distribution shifts provide a stringent test of robustness to unseen biological states.

**Fig. 7.**
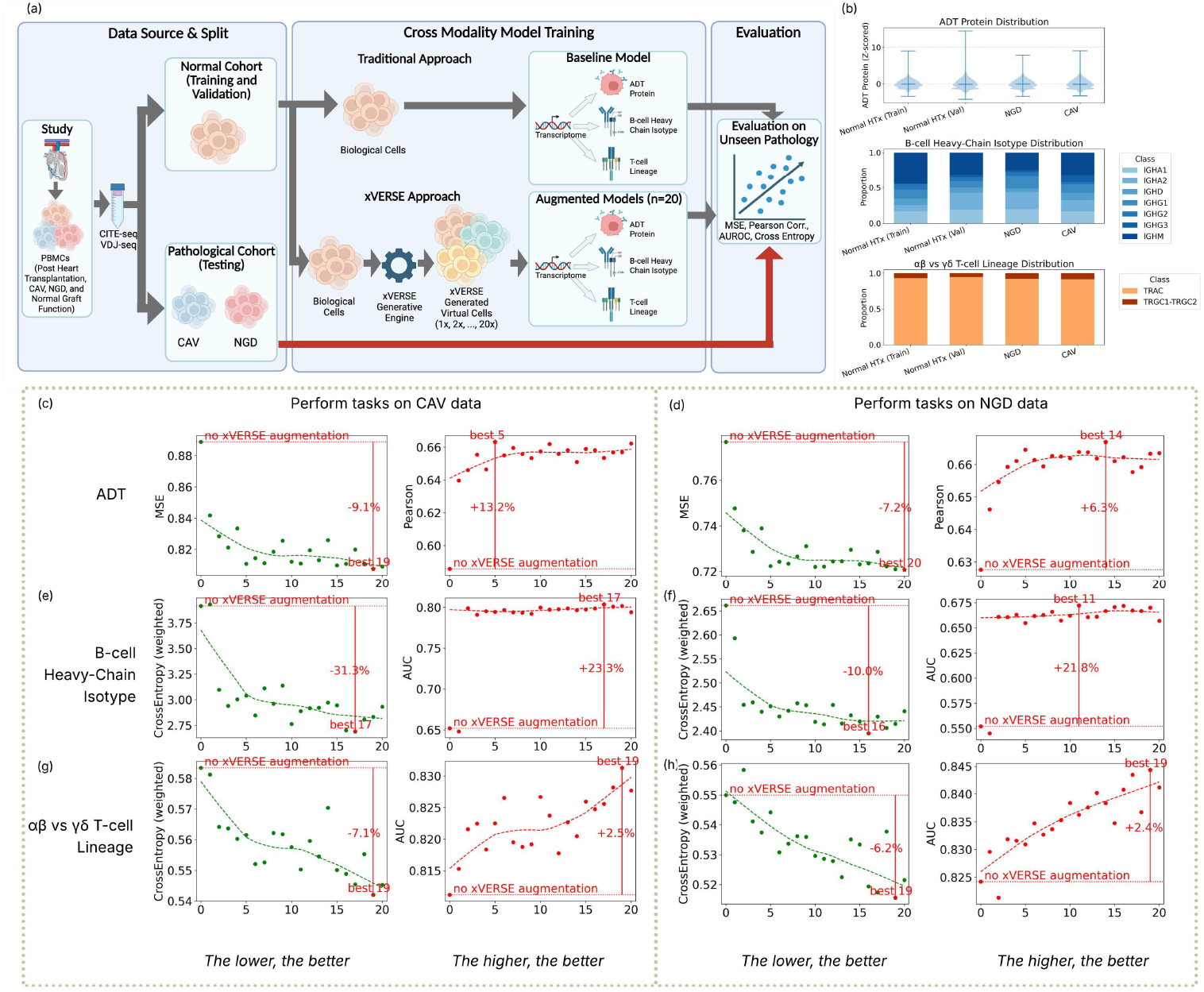
xVERSE-generated virtual cells facilitate robust cross-modality prediction and generalization to pathological states. **a**, Schematic overview of the cross-modality prediction benchmark. Models were trained to use cell transcriptomic profiles to predict ADT protein levels, B-cell heavy-chain isotype, and T-cell lineage using gene expression data. Training and validation were performed exclusively on *N* = 12 normal heart transplant (HTx) subjects (either biological cells alone or augmented with 1,2, 3, … and up to 20 virtual cells per biological cell). Then, the models (trained with biological cells only and xVERSE-augmented data) were tested on independent cohorts exhibiting Cardiac Allograft Vasculopathy (CAV) and Non-specific Graft Dysfunction (NGD). **b**, Distributional differences between training and evaluation cohorts. Violin plots show z-scored ADT marker distributions are broadly similar across cohorts with modest deviations, while stacked barplots reveal pronounced shifts in B-cell heavy-chain isotype usage and strong class imbalance in *αβ* versus *γδ* T-cell lineage labels (TRAC vs TRGC1–TRGC2) in NGD/CAV relative to the normal reference (training) cohort, highlighting the challenge of out-of-distribution generalization. **c**–**d**, Performance of ADT protein prediction. Comparison of Mean Squared Error (MSE) (**c**) and Pearson correlation (**d**) between models trained on biological cells only versus xVERSE-augmented data. Augmentation yields consistent error reduction and correlation improvement across both disease conditions. **e**–**f**, Evaluation of B-cell heavy-chain isotype classification. Scatter plots showing the reduction in CE and the increase in AUROC (**f**) tailored to isotype-specific signatures. The significant boost in AUROC highlights the model’s improved ability to capture rare isotype signals. **g**–**h**, Accuracy of T-cell lineage inference (*αβ* vs. *γδ*). Despite high baseline performance, augmentation further decreases CE (**g**) and improves classification AUROC (**h**), demonstrating xVERSE’s utility even in well-defined lineage tasks.

We compared two different training strategies: (i) training a baseline model using the original normal cohort and (ii) training a family of 20 models using 20 xVERSE-augmented normal cohorts; each augmented cohort contains *k* virtual cells per biological cell, with *k* ∈ { 1, 2, …, 20} . Remarkably, xVERSE-augmented models yielded substantial performance gains across all tasks and all pathological testing cohort compared to the baseline model (Fig. 7c–h). Specifically:

- **ADT Protein Prediction:** Augmentation reduced the Mean Squared Error (MSE) by 7.2%–9.1% and improved the Pearson correlation by 6.3%–13.2% (Fig. 7c–d), indicating that virtual cells facilitate a more accurate reconstruction of the proteomics landscape even across disease states. The augmentation improvements are stable as long as 5 or more virtual cells are synthesized per biological cell. Similar trends were observed in the other tasks.
- **B-cell Heavy-chain Isotype Classification:** We observed the most dramatic improvements in this task, where augmentation reduced Cross-Entropy (CE) by 10.0%–31.3% and increased the Area Under the Receiver Operating Characteristic Curve (AUROC) by over 20% (Fig. 7e–f). This suggests that virtual cells successfully capture and amplify subtle, isotype-specific transcriptomic signatures that are otherwise sparse in the biological data.
- **T-cell Lineage Prediction:** Even in tasks where baseline performance was already high, such as distinguishing *αβ* from *γδ* T-cell lineages, xVERSE augmentation further refined predictive accuracy, reducing CE by 6.2%–7.1% and improving AUROC by 2.5%–2.6% (Fig. 7g–h).

These results demonstrate that xVERSE-synthesized virtual cells provide a biologically coherent signal that reinforces model learning. By effectively expanding the training distribution, xVERSE enables downstream models to generalize more effectively to unseen pathological states, regardless of the baseline difficulty of the prediction task.

## 3 Discussion

We have presented xVERSE, a transcriptomics-native foundation model that fundamentally rethinks how we model cellular biology. While recent efforts have successfully adapted large language models (LLMs) to learn universal cell representations, these approaches often treat transcriptomic data as static tokens, overlooking the underlying probability distributions that govern gene expression. xVERSE bridges this gap by introducing a truly generative framework—one that does not merely embed cells into a latent space but learns to synthesize biologically faithful transcriptomic profiles from scratch.

This generative framework positions xVERSE as a powerful data augmentation engine that overcomes the physical constraints of experimental biology. Typically, increasing sample size or capturing rare cell states requires costly, labor-intensive experiments that are often infeasible with limited clinical tissue. xVERSE circumvents this bottleneck by synthesizing high-fidelity virtual cells that are statistically indistinguishable from real biological profiles. This allows researchers to computationally “perform experiments”—expanding small datasets and enriching rare populations—without generating new wet-lab data.

Crucially, xVERSE is designed to serve as a universal engine that can power any machine learning algorithm rooted in single-cell RNA-sequencing data. In the current era, where diverse single-cell machine learning algorithms are emerging rapidly, the potential of such a foundational engine is immense. We demonstrated this utility by augmenting training data with xVERSE-synthesized virtual cells, observing significant performance gains in cross-modality prediction tasks such as ADT protein abundance and immune receptor isotypes. This principle extends broadly to other machine learning challenges: xVERSE-synthesized virtual cohorts can be seamlessly integrated into existing analysis pipelines to enhance tasks ranging from automated cell-type annotation to trajectory inference and drug response prediction. This capability democratizes access to large-scale data, allowing researchers to train state-of-the-art models even when their own biological samples are scarce.

Intuitively, filling the latent manifold with these realistic virtual replicates rescues statistical power in low-n studies and enables the training of robust, generalizable machine learning models that would otherwise fail on sparse biological data alone.

Beyond cell-level tasks, xVERSE demonstrates exceptional flexibility in spatial transcriptomics through its ability to adapt to diverse gene panels. This capability is demonstrated in zero-shot and fine-tuned imputation, where xVERSE consistently outperforms specialized methods designed. By leveraging internalized universal priors to infer unmeasured genes from any given panel configuration, xVERSE effectively decouples biological discovery from hardware constraints, allowing researchers to extract maximal genomic insights from cost-effective, targeted experiments.

Finally, we demonstrated the effectiveness of the Gene2Cell score in deciphering the biological logic of the model. By quantifying the contribution of individual genes to specific cell representations, this metric effectively prioritizes biologically relevant features. Our results show that embeddings derived solely from high-scoring genes yield superior clustering fidelity, confirming that the model attends to the true drivers of cellular identity. Moreover, this scoring mechanism provides a data-driven strategy for spatial panel design, enabling the selection of minimal, high-information gene sets that retain full analytical power. These gene-level insights can be further integrated with pathway enrichment analyses and biological knowledge bases to enhance functional interpretation of cell states.

### Empirical Guidelines for Model Fine-tuning and Deployment

To facilitate the practical application of xVERSE, we provide empirical guidelines for fine-tuning and inference strategies tailored to different analytical scenarios.

#### Extreme Low-Data Regimes

For datasets with extremely limited cell numbers (e.g., rare clinical samples), we recommend using the full dataset for training without a validation split. The model should be fine-tuned until the loss on the training set stabilizes. In this regime, maximizing the utilization of every available cell is critical for capturing the latent manifold of rare populations.

#### Standard Analysis Tasks

For larger datasets where the goal is data augmentation or general analysis, we advise setting aside 20-30% of the data for validation. Training should proceed until the loss on the validation set stabilizes. During inference/generation, we recommend using the batch-conditioned Poisson parameters (incorporating batch information) to synthesize virtual cells that faithfully reflect the technical characteristics of the target dataset, ensuring seamless integration with the original data.

#### Imputation Tasks

For imputation in spatial transcriptomics, performance can be enhanced by co-training with a paired single-cell reference dataset matched for tissue and disease state. We recommend setting aside 20-30% of the data for validation. Training should proceed until the loss on the validation set stabilizes. Crucially, for the final imputation step, we suggest using the batch-corrected Poisson parameters (removing batch information) to recover the underlying biological expression profile free from technical artifacts.

### Limitations

Our study has limitations. First, xVERSE is explicitly designed to model the high-dispersion count distributions characteristic of droplet-based single-cell and imaging-based spatial technologies. Consequently, its adaptability to non-UMI protocols like Smart-seq remains limited, as they are prone to PCR amplification bias and follow noise distributions that diverge from the discrete count models optimized for UMI-based sequencing. Second, as multi-omics technologies mature, future iterations of xVERSE will need to incorporate chromatin accessibility (scATAC-seq) and protein abundance (CITE-seq) into its latent cell space to provide a truly holistic view of cell state.

In conclusion, xVERSE stands as the first generative foundation model natively designed for transcriptomics, fundamentally expanding the scope of single-cell analysis. By enabling researchers to computationally extend their experiments through virtual synthesis, xVERSE establishes a new paradigm for data-driven discovery in the era of large-scale genomics.

## STAR Methods

### Training Data Collection and Filtering

We collected a comprehensive training dataset comprising all publicly available singlecell RNA-sequencing data from the CellxGene Census (census version is “2025-01-30”) and all spatial transcriptomics datasets from the official repositories of 10x Genomics (Xenium), NanoString (CosMx), and Vizgen (MERFISH).

To ensure data quality, we applied a rigorous two-stage filtering strategy. First, for each dataset, we removed low-quality cells and potential doublets using adaptive thresholds based on the distribution of detected genes and total UMI counts. Specifically, we excluded cells where the number of detected genes exceeded the median plus three standard deviations, and where the total UMI count exceeded the minimum of 20,000 or the median plus three standard deviations.

Second, during data loading, we applied an additional filter to exclude cells with extreme expression values, removing any cell where the maximum UMI count for a single gene exceeded 1000. This step ensures the removal of technical artifacts while retaining the vast majority of biologically valid cells.

### Standardization of Cell Type Annotations

While the CellxGene Census provides extensive cell type annotations, these labels lack uniformity and often exhibit hierarchical dependencies (e.g., “fast muscle cell” is a subtype of “muscle cell”), rendering them unsuitable for direct large-scale training. To resolve this, we utilized a custom prompt (see Suppl. Note) to map the original Cell Ontology annotations to a set of 29 broad, standardized cell type categories defined in our study. Cell types that could not be unambiguously mapped or fell outside these major lineages were classified as “Other/Unknown”.

### xVERSE Training Architecture

#### Input Gene Expression Preprocessing

For an individual cell *i*, we define the input gene expression profile as a vector ***x***_*i*_ ∈ ℝ^*G*^, where *G* = 17, 999 represents the gene universe, defined as genes with non-zero expression in at least 50% of the collected samples.

Raw counts are processed into a dense value vector ***v***_*i*_ where measured entries are log-transformed (*v*_*ij*_ = log(1+*x*_*ij*_)) and unmeasured entries are zero-padded. Each cell *i* is associated with a binary measurement mask vector ***m***_*i*_ ∈ {0, 1}^*G*^, where *m*_*ij*_ = 1 indicates gene *j* was measured in cell *i* or not. This mask helps to differentiate genes are measured but not expressed or not measured. In addition, the following labels are used for cell *i*: a tissue identifier *t*_*i*_, and during pre-training, a sample identifier *s*_*i*_ and cell type label *c*_*i*_. Cells without cell-type labels will have *c*_*i*_ = *−* 1.

To simulate the sparsity and varying gene coverage inherent in spatial transcriptomics data, we employed a dynamic random masking strategy during training. For cells with low coverage (*<* 1000 detected genes), we randomly masked between 10 genes and 20% of the detected genes. For cells with higher coverage, we applied a stochastic masking approach: 90% of the time, we randomly masked a fraction of detected genes (10–30% with *p* = 0.2, 30–50% with *p* = 0.7, or 50–75% with *p* = 0.1). The remaining 10% of the time, we simulated panel-based acquisition by applying predefined gene panels, further augmented by randomly dropping 10–100 genes or adding back 50–3000 genes to mimic technical noise and panel variation.

### The Cell and Gene Encoder

xVERSE encoder derives gene representations via adaptively learned gene representations, and then transform them to cell embeddings. The details are as follows.

Each gene has a **global gene embedding *E***_gene_[*j*] ∈ ℝ^*D*^, which is adaptively learned during training.

To account for the different gene expression patterns across tissues, we use a tissue-specific gene embedding lookup table ***T*** _tissue_, where each gene *j* in tissue *t*_*i*_ is associated with a scalar value ***T*** _tissue_[*t*_*i*_, *j*], which is adaptively learned during training.

For cell *i*, we construct a composite input feature vector 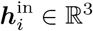 by concatenating the processed expression value *v*_*i*_, the mask status *m*_*i*_, and ***T*** _tissue_ [*t*_*i*_, *] . Then, 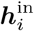 are projected to a value via a Multilayer Perceptron (MLP) (Φ_val_ : ℝ^3^ → ℝ^1^). The weights in the MLP are initial unnormalized weights, which are then processed by a mixing block (Φ_mix_ : ℝ^*G*^ → ℝ^*G*^) that exchanges information across genes to refine the weights. Let 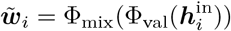 be the refined weights.

To eliminate the contribution of unmeasured genes to the embedding, we mask the unmeasured and randomly masked genes.

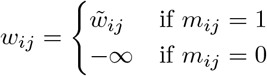

Then, we perform a softmax normalization ***α***_*i*_ = Softmax(***w***_*i*_) to obtain the **Gene2Cell** score for cell *i*.

The preliminary embedding for cell *i* is computed as the weighted sum of learnable gene embeddings ***E***_gene_ *∈* ℝ^*G×D*^:

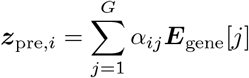

To adapt to varying gene panels, ***z***_pre,*i*_ is modulated by a Feature-wise Linear Modulation (FiLM) layer conditioned on the specific mask pattern ***m***_*i*_, followed by an output projection to yield the final **cell embedding *z***_bio,*i*_ *∈* ℝ^*D*^.

### Cell-Gene Poisson Parameter Decoder

The decoder reconstructs two Poisson parameters *µ*_bio,*ij*_ and *µ*_sid,*ij*_ for cell *i* and gene *j* based on the latent cell embedding ***z***_*bio,i*_, gene embedding ***E***_*gene*_[*j*], tissue-gene bias ***B***_tissue_[*t*_*i*_, *j*], and a sample embedding ***e***_*si*_ corresponding to the cell’s batch ID. These two Poisson parameters are used to reconstruct the gene expression profile of cell *I* with and without sample-specific batch effects, respectively.

For each cell *i* and gene *j*, we assemble a query vector by concatenating the cell embedding ***z***_bio,*i*_ with the corresponding gene embedding ***E***_*gene*_[*j*]. This query vector is passed through a shared gene decoder MLP to predict a logit parameter *y*ŷ_*ij*_, which is then adjusted by adding a separate learnable tissue-gene bias ***B***_tissue_[*t*_*i*_, *j*]. A separate MLP predicts the total library size *l*_*i*_ from ***z***_bio,*i*_, followed by a Softplus activation to ensure positivity. The mean parameter of the Poisson distribution is given by *µ*_bio,*ij*_ = *l*_*i*_ Softmax_*j*_(ŷ_*ij*_ + ***B***_tissue_[*t*_*i*_, *j*]).

Further, to incorporate sample-specific contextual information, a FiLM layer can be added to modulate cell embeddings.

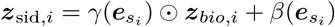

Following the same architecture mentioned above, we also decode *µ*_sid,*ij*_.

### Loss Functions and Optimization

The model is trained end-to-end using a composite objective function that balances between generative reconstruction accuracy and biological signal retention.

- Poisson Reconstruction Loss: We employ the Poisson negative log-likelihood loss to ensure accurate reconstruction of cellular transcriptomic profiles. This is computed for both Poisson parameters ***µ***_bio_ and ***µ***_sid_, strictly over the set of measured genes. For a single cell *i*, the loss is defined as:

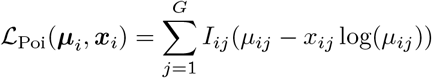

where *I*_*ij*_ is the binary indicator of whether gene *j* was physically measured in cell *i* (i.e., the panel genes, excluding the random masking applied during training).
- Adversarial Batch Correction: To disentangle biological signals from technical artifacts, we employ a conditional adversarial training strategy. We introduce a sample discriminator *f*_adv_ that attempts to predict the sample ID *s*_*i*_ from the biological latent ***z***_bio,*i*_. To enforce batch invariance, we apply a Gradient Reversal Layer (GRL) to the input of *f*_adv_, reversing the gradient sign during backpropagation. To prevent the removal of biological variation (e.g., cell type composition differences across batches), we perform this adversarial training within each cell type. The discriminator is trained using a cross-entropy loss ℒ_batch_ computed via a conditional batch strategy: for each cell type, we balance the sample IDs to the median count via resampling (downsampling or upsampling) before computing the loss on this balanced subset.
- Cell Type Classification Loss: To supervise the biological latent space ***z***_bio_ to capture cell type-specific information, we employ a classifier *f*_cls_ that predicts cell type logits from ***z***_bio,*i*_. The model is trained with a standard cross-entropy loss ℒ _celltype_ using label smoothing (*ϵ* = 0.05) to encourage robust cluster separation. The final objective function combines these components, weighted by hyperparameters to balance their contributions:

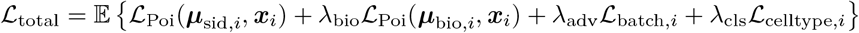

where *λ*_*bio*_, *λ*_*adv*_, and *λ*_*cls*_ are hyperparameters controlling the strength of the biological reconstruction, adversarial batch removal, and cell type classification tasks, respectively.

### Computation Resources Used for Training

Pretraining was conducted on a high-performance computing (HPC) node equipped with NVIDIA H200 GPUs, with each job allocating 6 H200 GPUs, 20 CPU cores, and 1450 GB of system memory.

### xVERSE Fine-Tuning Architecture

During fine-tuning, due to the limited sample size of the dataset, we disabled the Adversarial Batch Correction module and the Cell Type Classification module. The model was optimized solely based on the Poisson reconstruction loss to adapt the biological representations and the decoder to the specific dataset characteristics.

Some fine-tuning tasks were performed on an HPC node equipped with 1 A6000 GPU, 20 CPU cores, and 200 GB of system memory; and others were performed on an HPC GPU node equipped with 1 H200 GPU, 20 CPU cores, and 200 GB of system memory.

### xVERSE Performance Benchmark and Evaluation

#### Benchmarking Universal Cell Representations

To rigorously assess the robustness of xVERSE, we utilized two distinct datasets: the Single-Cell Atlas of Human Pediatric Liver and a multiome dataset comprising healthy and ALS human motor cortex and spinal cord samples. We compared xVERSE against three state-of-the-art foundation models (scGPT, Nicheformer, and Geneformer) and the specialized integration method, Harmony. Consistent with a “universal” application, xVERSE and the foundation models were evaluated via zero-shot inference without fine-tuning, while Harmony was used to explicitly integrate batches. We simulated diverse experimental scenarios by assessing performance across three input gene panels: (1) the whole transcriptome, (2) the Xenium Prime Panel (∼ 5,000 genes), and (3) tissue-specific Xenium panels (377 genes for liver and 266 genes for brain). Cells were projected to the latent space of different models.

To quantify the quality of the latent space, we applied the `scib.metrics.silhouette` function in the `scib`[86] package to compute the Average Silhouette Width (ASW) based on cell type identity. The silhouette width for a single cell *i* is defined as 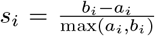, where *a*_*i*_ is the mean distance between cell *i* and all other cells in the same cluster (cell type), and *b*_*i*_ is the mean distance between cell *i* and all cells in the nearest distinct cluster. The raw ASW is the average of *s*_*i*_ across all cells, ranging from -1 to 1. To facilitate interpretation, the final score is normalized to the range [0, 1] using the formula *ASW* = (*ASW*_*raw*_ + 1)*/*2, where 0 indicates random assignment and 1 indicates perfect separation of cell types.

For fair comparison, all models reported in Fig. 2 (xVERSE and other foundation models) were inferenced under a configuration allocating 3 H200 GPUs, 20 CPU cores, and 200 GB of system memory.

### Evaluating the Biological Relevance of Gene2Cell Scores

To determine whether the learned Gene2Cell scores capture biologically meaningful signals, we fine-tuned xVERSE on 2 cohorts: five human breast single-cell samples (PM-A to PM-E) and four CosMx lung samples (Lung5_Rep1, Lung6, Lung9_Rep1, Lung13F), respectively. Post-fine-tuning, we calculated the average Gene2Cell score for every gene across all cells in each dataset, ranked the genes by these scores, and stratified them into different quantile groups.

First, using the human breast single-cell samples,we assessed the representational capacity of these strata to determine if high-scoring genes alone can recapitulate the full transcriptomic state. We generated cell embeddings using only the genes within each quartile group (0–25%, 25–50%, 50–75%, and 75–100%) and evaluated their quality using the Average Silhouette Width (ASW), comparing them against embeddings derived from the full transcriptome.

Second, using the CosMx lung samples, we evaluated the predictive utility of these genes by testing their ability to impute 50 held-out highly variable genes (HVGs) masked. We used the gene sets from each quantile group (top 50% vs bottom 50%, top 33% vs mid 34% vs bottom 33%, top 25% vs 25%–5s0% vs 50%–75% vs bottom 25%) as inputs for the imputation task (details in Section 3).

### Evaluating Virtual Cell Synthesis Ability and Fidelity

We fine-tuned xVERSE on single-cell human breast tissue datasets (samples PM-A to PM-E). Raw gene expression matrices were processed to retain high-quality cells based on standard quality control (QC) metrics: cells with fewer than 200 detected genes, fewer than 200 total counts, or greater than 20% mitochondrial content were excluded. Genes with zero counts across all cells were removed. The filtered data was used as input for fine-tuning.

We fine-tuned the pre-trained xVERSE model on all samples together. The dataset was split into 90% training and 10% validation sets. The model was trained using the Adam[87] optimizer with an initial learning rate of 1 *×* 10^*−*3^, which was dynamically reduced by a factor of 0.8 if the validation loss did not improve for 5 epochs. Training proceeded for a maximum of 300 epochs with an early stopping mechanism triggered after 20 epochs of no improvement. The fine-tuned model decoded sample-specific Poisson parameter vector for each cell. A virtual cell is a random draw from the Poisson distribution with the sample-specific Poisson parameter vector. Multiple virtual samples can be synthesized from a given biological sample.

Virtual cell’s fidelity was evaluated by (a) comparing the marginal distributions of UMI counts for individual genes across virtual cells in each virtual sample versus biological cells; (b) gene-wise Pearson correlation between the observed gene expressions and the model’s output Poisson parameter vector for the top *K* HVGs (varying *K* from 50 to 1000); and (c) Average Silhouette Width (ASW) (using `scib.metrics.silhouette` function in the `scib` package) based on cell type identity using embeddings following standard Scanpy[88] 1.10.4 pipeline.

In addition, we trained a discriminative classifier to distinguish biological cells from virtual cells. We constructed a balanced dataset of biological cells and virtual cells and split it into 70% training and 30% testing sets. A logistic regression classifier was trained to distinguish biological cells from virtual cells.

We benchmarked xVERSE against scVI[89], a non-foundation-model based but widely used generative model capable of synthesizing virtual cells. We trained the model using the `scvi.model.SCVI` class with default hyperparameters and early stopping enabled. To synthesize virtual cells, we employed the `posterior_predictive_sample` function, generating 3 virtual replicates for each biological cell.

### Benchmarking Gene Imputation Capability

We utilized a human lung dataset profiled with the CosMx SMI platform (4 samples: Lung5_Rep1, Lung6, Lung9_Rep1, Lung13). For each sample, we identified the top 50 HVGs using the `scanpy.pp.highly_variable_genes` function with `flavor=‘seurat_v3’`, and masked them during model fine-tuning and inference. These 50 HVGs are the “test set” for imputation evaluation.

We applied the following four imputation strategies, and compared their performances.

- **xVERSE (zero-shot)**: We directly applied the pre-trained xVERSE model to the spatial data without fine-tuning. The output biological Poisson parameter vector (`mu_bio`) for each cell was used as the imputed gene expression values.
- **xVERSE (fine-tuned)**: We followed the standard xVERSE fine-tuning protocol to fine-tune the model on a composite dataset consisting of the 4 spatial samples (with masked HVGs) and 5 matched single-cell reference datasets. The output biological Poisson parameter vector (`mu_bio`) for each cell was used as the imputed gene expression values.
- **SpaGE**: SpaGE is a domain adaptation method that aligns single-cell and spatial data using the PRECISE algorithm, followed by k-nearest-neighbor (kNN) regression to predict unmeasured gene expression. Since this methods requires a reference single-cell dataset, we used five different NSCLC single-cell samples as the reference. We used the `SpaGE` function with `n_neighbors=50`.
- **gimVI**: gimVI is a generative model for integrating spatial and single-cell data. We used the `scvi.external.GIMVI` class with default parameters and `max_epochs=20`, using 5 different NSCLC single-cell samples as the reference.

### Evaluating whether xVERSE-generated virtual cells can enhance standard single-cell analysis in data-poor regimes

To evaluate xVERSE’s capability to enhance analysis in data-poor regimes, we designed two experimental scenarios using the multimodal PBMC dataset. This study profiled human PBMCs using both 3’ and 5’ 10x Genomics chemistries, allowing us to evaluate performance across different library preparation protocols.

First, we focused on the 3’ dataset, and considered four challenging clustering tasks involving two cell types: (1) 5 ILC vs. 55 NK cells, (2) 6 Plasmablasts vs. 54 B memory cells, (3) 10 CD4 Proliferating vs. 50 HSPC cells, and (4) 4 Erythrocytes vs. 56 Platelets. We fine-tuned xVERSE on each small dataset for 25 epochs. For inference, we generated 5 virtual cells for each biological cell. We then used the `sc.tl.leiden`[81] function in the scanpy package to perform Leiden clustering (resolution 0.5) on the virtual cells and assigned a consensus label to each original cell via majority voting. We benchmarked this “xVERSE Majority Vote” approach against a standard analysis pipeline (Leiden clustering at resolutions 0.5, 1.0, and 2.0) and quantified performance using the Adjusted Rand Index (ARI) and Normalized Mutual Information (NMI).

Next, we focused on the 5’ dataset, and performed a cell-type-stratefied sub-sampling of the full dataset to retain only 1% of the total cells per cell type. We fine-tuned xVERSE on this 1% subsample for 1500 epochs and generated 100 virtual cells for each biological cell. Then, we performed Wilcoxon rank-sum test using the `sc.tl.rank_genes_groups` function in the `scanpy` package to identify differentially expressed genes (DEGs) for each cell type. This approach was called “xVERSE augmentation”. We benchmarked this approach against a “baseline oversampling” approach (naively copying each cell 100 times). We evaluated performance by comparing the Precision and Recall of identified DEGs against a Ground Truth derived from the full, original dataset (p-value *<* 0.05). We computed these metrics across a wide range of p-value thresholds (from 10^*−*8^ to 0.05) to generate Precision-Recall curves.

### Evaluating xVERSE as a data augmentation engine for downstream machine learning model training

We performed a task on a multi-modality (CITE-seq and VDJ-seq) multi-cohort heart transplant dataset with three cohorts: normal controls with no complications after transplant, patients with Non-specific Graft Dysfunction (NGD), and patients with Cardiac Allograft Vasculopathy (CAV).

The goal is to use the gene transcriptomics profile to predict (1) Antibody-Derived Tags (ADT protein levels; measured by CITE-seq) and (2) B-cell heavy chain isotypes (measured by VDJ-seq) and (3) T-cell lineages (*αβ* versus *γδ*; measured by VDJ-seq). For each task, we performed a generalization benchmark where models were trained exclusively on samples from normal controls and evaluated on NGD and CAV samples.

The dataset was split for training, validation, and testing as follows:

- Training Set: 8 healthy control samples (CONTROL1_1 to CONTROL4_2).
- Validation Set: 4 healthy control samples (CONTROL5_1 to CONTROL6_2).
- Test Set 1: 6 NGD samples.
- Test Set 2: 6 CAV samples.

We trained two models with and without xVERSE data augmentation.

- Baseline model: Only biological cells from the training set are used to train the prediction models.
- xVERSE-augmented model: We fine-tuned xVERSE on the training set using the Adam optimizer with a learning rate of 1×10^*−*4^ for a maximum of 50 epochs, with early stopping triggered after 5 epochs of no improvement on the validation set. Then, we synthesized 1, 2, up to 20 virtual cells for each biological cell in the training set using the fine-tuned xVERSE model. Then, we used the augmented training set (with both biological cells and virtual cells) to train the prediction models.

All prediction models were trained using a Multi-Layer Perceptron (MLP) architecture. The MLP consists of two hidden layers with 128 units each, followed by ReLU activation and dropout (rate=0.2). The output layer size corresponds to the number of targets (ADT proteins or V(D)J classes). Models were trained for 100 epochs with the Adam optimizer (learning rate 1 *×* 10^*−*4^) and early stopping (patience 10).

## Data and Code Availability

### Data Availability

The datasets used in this study are publicly available from the following repositories:

- **Training Data:** Single-cell RNA-sequencing data was obtained from the Cellx-Gene Census (version 2025-01-30). Spatial transcriptomics training data includes all public datasets available from the official websites of 10x Xenium, NanoString CosMx, and Vizgen MERFISH prior to June 2025.
- **Universal Representation Benchmarks (Fig. 2):** The Human Pediatric Liver atlas is available from CellxGene (Collection ID: ff69f0ee-fef6-4895-9f48-6c64a68c8289, Dataset ID: 10cc50a0-af80-4fa1-b668-893dd5c0113a). The ALS motor cortex multiome dataset is available from CellxGene (Collection ID: 0986e4cd-7a58-405d-9b91-4b199bb4124e, Dataset ID: 0ab54d91-066c-4223-a9ea-6a3b0d1adef4).
- **Virtual Cell Synthesis (Fig. 3, 4):** The human breast single-cell dataset (PM-A to PM-E) is available under GEO accession GSE180878.
- **Spatial Imputation (Fig. 3, 5):** The human lung NSCLC CosMx SMI dataset (Lung5_Rep1, Lung6, Lung9_Rep1, Lung13) is available from the NanoString website (https://nanostring.com/products/cosmx-spatial-molecular-imager/ffpe-dataset/nsclc-ffpe-dataset/).
- **Data-Poor Regime Analysis (Fig. 6):** The multimodal PBMC dataset (3’ and 5’ kits) is available under GEO accession GSE164378.
- **Cross-Modality Prediction (Fig. 7):** The heart transplant CITE-seq and VDJ dataset is available under GEO accession GSE291290.

## Code Availability

The source code for xVERSE, including model training, inference, and analysis scripts, is available on GitHub at https://github.com/jichunxie/xVERSE_code. Custom scripts for reproducing the figures in this manuscript are also provided in the repository.

## Declaration of generative AI and AI-assisted technologies in the manuscript preparation process

During the preparation of this work the author(s) used Google Gemini 2.5, Google Gemini 3, and OpenAI Codex in order to format the code. The author(s) also used Google Gemini 3 to polish the writing of this manuscript. After using this tool/service, the author(s) reviewed and edited the content as needed and take(s) full responsibility for the content of the published article.

## Acknowledgments

We would like to thank Cliburn Chan, John Hickey, and Andrew Nixon for their helpful suggestions and feedback. We thank Yutong Cheng for designing the logo of xVERSE. X. Jiang’s research was supported by the NIH award R01HG012555 and Duke University. J. Xie’s research was supported by the NIH awards R01HG012555, U54AG075936, U01AI186999, and Duke University. The findings and conclusions presented in this paper are those of the author(s) and do not necessarily reflect the views of the NIH.

